# Allele-specific CRISPR/Cas9 editing inactivates a single nucleotide variant associated with collagen VI muscular dystrophy

**DOI:** 10.1101/2024.03.22.586265

**Authors:** Véronique Bolduc, Katherine Sizov, Astrid Brull, Eric Esposito, Grace S. Chen, Prech Uapinyoying, Apurva Sarathy, Kory Johnson, Carsten G. Bönnemann

## Abstract

The application of allele-specific gene editing tools can expand the therapeutic options for dominant genetic conditions, either via gene correction or via allelic gene inactivation in situations where haploinsufficiency is tolerated. Here, we used allele-targeted CRISPR/Cas9 guide RNAs (gRNAs) to introduce inactivating frameshifting indels at a single nucleotide variant in the *COL6A1* gene (c.868G>A; G290R), a variant that acts as dominant negative and that is associated with a severe form of congenital muscular dystrophy. We expressed spCas9 along with allele-targeted gRNAs, without providing a repair template, in primary fibroblasts derived from four patients and one control subject. Amplicon deep-sequencing for two gRNAs tested showed that single nucleotide deletions accounted for the majority of indels introduced. While activity of the two gRNAs was greater at the G290R allele, both gRNAs were also active at the wild-type allele. To enhance allele-selectivity, we introduced deliberate additional mismatches to one gRNA. One of these optimized gRNAs showed minimal activity at the WT allele, while generating productive edits and improving collagen VI matrix in cultured patient fibroblasts. This study strengthens the potential of gene editing to treat dominant-negative disorders, but also underscores the challenges in achieving allele selectivity with gRNAs.

## Introduction

Collagen VI is a secreted, multimeric protein abundant in the connective tissues of skeletal muscle.^1,2^ The main *COL6* genes, *COL6A1*, *COL6A2*, and *COL6A3*, are expressed by interstitial muscle fibroblasts and not by myofibers.^3,4^ They encode three proteins, termed, respectively, collagen α1, α2, and α3(VI) chains, each structured with a collagenous central triple helical domain composed of Gly-X-Y repeats.^5^ Owing to the small glycine in these repeats, the three alpha chains can tightly intertwine to form a monomer, composed of one of each of the chains, that further self-assemble into dimers and tetramers.^6,7^ Tetramers are secreted and polymerize end-to-end into a microfibrillar network that constitutes the collagen VI matrix^6^ and that contacts with myofibers.^3^

Integrity of the triple helical domains of collagen α1, α2, and α3(VI) chains is thus essential for proper collagen VI production. Dominantly acting variants account for over 65% of all *COL6* pathogenic variants that cause collagen VI-related muscular dystrophies (COL6-RD),^8^ and they cluster at the amino-termini of the triple helical domains.^2^ Initial coiling of the alpha chains, to form monomers, proceeds from the carboxy-towards the amino-termini of the triple helical domains. As a result, pathogenic variants at the amino-termini allow the alpha chains harboring them to be incorporated, rather than excluded, into monomers, and to further propagate to dimer and tetramer stages.^9^ For this reason, these “assembly-competent” variants exert strong dominant-negative effects. One type of variant that is frequently reported is glycine substitutions of the first position of the Gly-X-Y repeat.^2,10^ Typically, such glycine substitutions are not tolerated and cause structural changes to tetramers that appear “kinked” on electron microscopy and that exhibit reduced ability to polymerize in the matrix.^9,11^ By mechanisms not yet fully elucidated, dysfunctional collagen VI production leads to skeletal muscle weakness, progressive joint contractures, and respiratory failure, which can be fatal.^1,12^

There are currently no treatments for COL6-RD. Given that *COL6A1*, *COL6A2* and *COL6A3* genes are haplosufficient, i.e., losing or inactivating one copy of either of these genes does not result in a clinical phenotype,^12,13^ we and others have postulated that allele-specific silencing would be a promising therapeutic strategy for dominant-negative variants causing COL6-RD. This can be achieved with antisense technologies, such as small interfering RNAs or gapmer antisense oligonucleotides,^14^ adapted to be allele-specific. COL6-RD primary cultures treated with these molecules provided evidence that specifically knocking down the pathogenic transcripts attenuates the dominant-negative effect and improves collagen VI matrix deposition;^15–18^ however, complete and sustained mutant allele knockdown with stringent allele selectivity has yet to be demonstrated in preclinical animal models of COL6-RD.

Gene editing via the Clustered Regularly Interspersed Short Palindromic Repeats (CRISPR)/Cas9 system is also adaptable to achieve allele-specific gene inactivation.^19^ Engineered CRISPR/Cas9 is a two-component system that includes expression of a nuclease (e.g., *Streptococcus pyogenes* Cas9, SpCas9) concomitantly with a single guide RNA (gRNA) to induce a double-stranded break at desired genomic locations.^20^ Using a gRNA sequence complementary to a single allele enables allele-specific targeting, even for single nucleotide changes.^19^ Creation of double-stranded breaks then activates the cellular DNA repair pathways, which, in absence of template, are conducted primarily by the canonical non-homologous end joining (NHEJ) pathway, or by the microhomology-mediated end joining (MMEJ) pathway (also called alternative end-joining) (reviewed in ^21,22^). Both NHEJ and MMEJ are error-prone and readily introduce indels during the repair process following a Cas9-induced double-stranded break. The DNA modifications, if frameshifting, can permanently inactivate expression of the gene in which they were introduced.

Allele-specific CRISPR/Cas9-induced gene inactivation has been tested in various cellular and animal models of dominant diseases, with variable degrees of Cas9 on-target activity and allele discrimination.^23–29^ Here, we applied this precision medicine to a recurrent glycine substitution in *COL6A1* (G290R).^9,10^ We show that the indel profiles favor gene inactivation, and that the gRNA design can be optimized for enhanced allele selectivity with the addition of base mismatches.

## Results

### CRISPR/Cas9 guide RNAs selected for the *COL6A1* single nucleotide variant (c.868G>A; G290R) preferentially introduce frameshifting edits at the variant allele

At the *COL6A1* c.868G>A locus, we selected two gRNAs (gRNA-A and gRNA-B; Figures 1A, S1A,B) in which the c.868 nucleotide was located at position 1 of the protospacer, i.e. at a 1-nucleotide distance relative to the NGG protospacer adjacent motif (PAM). We prepared plasmid constructs, each expressing an SpCas9-GFP fusion protein concomitantly with either gRNA-A or gRNA-B, or without gRNA (No gRNA; Figure S1C). To test our gene editing strategy, we nucleofected the constructs into primary dermal fibroblast cells obtained from four independent patients carrying the *COL6A1* c.868G>A (G290R) variant (Pt1 to Pt4), and from one non-neuromuscular disease patient as a control (Ctrl). Dermal-derived fibroblasts express collagen VI at high levels and are therefore a good surrogate for the muscle interstitial fibroblasts.^30^ GFP-enriched cell populations, collected 48h after nucleofection, were used for all further analyses (Figure S1C).

**Figure 1.**
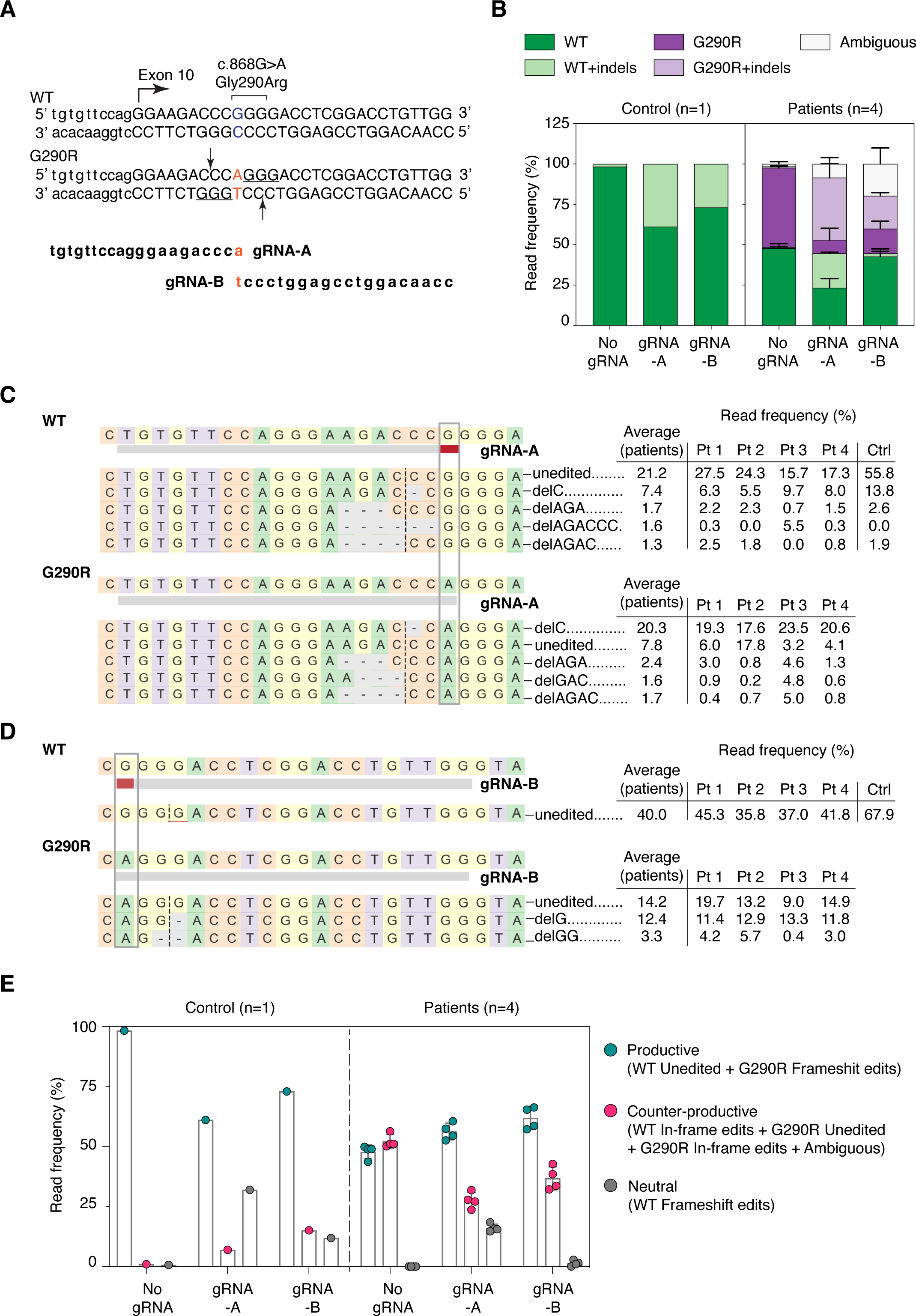
Two selected gRNAs introduce frameshifting edits preferentially, but not specifically, at the *COL6A1* c.868G>A (G290R) variant allele. (**A**) Two guide RNAs (gRNA-A and gRNA-B) were designed to specifically anneal to the *COL6A1* c.868G>A; G290R (marked in red) variant. The PAM sequences utilized (underlined) and the Cas9 cleavage sites (arrows) are shown. **(B)** Targeted re-sequencing (Illumina Mi-Seq) of the *COL6A1* c.868G>A locus was performed in four patient and one control primary cells after gene editing with gRNA-A, gRNA-B, or without gRNA (No gRNA). Sequencing reads were analyzed by Crispresso2 and assigned to either the native or the variant allele (WT or G290R) and called whether they contain indels (+ indels) or not. Reads that couldn’t be assigned to either allele were termed ambiguous. Histogram plots distribution of the read frequencies, expressed as percentage of total reads for each sample. For patient samples, bars represent the average of four individuals, ± standard deviation. (**C**) Alignment of the most frequent reads (reaching a read frequency of at least 1.5% in either the patient or the control samples), for gRNA-A. A single frameshifting edit (delC) was the most frequent edit at the G290R allele in all 4 individuals, surpassing unedited G290R reads, as shown by the individual read frequencies reported. DelC was the second most frequent read at the WT allele, after unedited WT. **(D)** Alignment of the most frequent reads, for gRNA-B. Similarly to gRNA-A, a single frameshifting edit (delG) was the most frequent repair at the G290R allele, after unedited G290R reads, in all 4 individuals. (**E**) Reads were compiled according to the functional outcome, whether they were 1) productive (unedited WT reads, or G290R reads containing a frameshifting edit), 2) damaging (that likely produces a dominant-negative product, such as unedited G290R reads, WT or G290R reads containing in-frame edits, or ambiguous reads that encompass the splice site), or 3) neutral (WT reads that include frameshifting edits). n=4 patient lines ± standard deviation.

We performed targeted deep-sequencing of the locus of interest, to measure the precise editing frequencies (Figure S1C). The total number of reads was tallied for each sample (average of 188,607 reads per sample; Figure S2A) and served as denominators to calculate read frequencies. Both gRNAs efficiently introduced indels at the c.868G>A (G290R) allele in patient cells (38.4% and 20.3% of total reads were G290R allele modified with indels, for gRNA-A and gRNA-B, respectively, vs 0.8% for No gRNA; Figure 1B). While they were active at the G290R allele, both gRNAs also modified the native (wild-type, WT) allele in patient cells (21.2% and 1.9% of total reads, for gRNA-A and gRNA-B, respectively), and in the control sample (38.9% and 27.0% of total reads, for gRNA-A and gRNA-B, respectively). With gRNA-B, deletions frequently encompassed the c.868 nucleotide site, such that a larger proportion of reads were ambiguous (19.8% of total reads; Figure 1B). It is likely that a fraction of these represent modified WT reads and that the 1.9% read frequency observed in this sample is underestimated. The 27.0% of modified reads in the control sample suggests that gRNA-B does not effectively discriminate between the two alleles. Overall, both gRNAs were more active at the G290R allele compared to the WT allele, but they did not fully discriminate the G290R from the WT alleles.

We next examined the indel edits generated by either gRNAs. For gRNA-A, in patient cells, deletions were observed far more frequently than insertions (54.7% of total reads – G290R and WT alleles combined – vs 4.0% of total reads, respectively; Figure S2B). Of these, 1-bp deletions were the most frequent (33.0% of total reads, G290R and WT alleles combined; Figure S2C). A single edit, deletion of a cytosine at the cleavage site (c.865delC), accounted for most of the modified reads (27.7% of total reads, for G290R and WT alleles combined; Figure 1C and S2D). Looking at the G290R allele only, this edit represented nearly half of the reads (43.4% of G290R reads, n=4 patient cells). Preference for this modification was observed consistently in the four patient cell lines tested (Figure 1C and S2D), as well as in the control cell line (13.8% of total reads; Figure 1C and S2D). Similarly, for gRNA-B, the most frequent edits were deletions (19.9% of total reads, for G290R and WT alleles combined), and more precisely, one-bp deletions (13.6% of total reads, for G290R and WT alleles combined; Figure S2B,C). The ambiguous reads were also modified with deletions but were not tallied in these analyses since they could not be attributed to either allele. For gRNA-B, the single most frequent edit was c.871delG (12.4% of total reads for G290R and WT alleles combined), and was also observed as the preferential indel in the four patient cell lines tested and in the control cell line (Figure 1D and S2D).

While the preferential edits (c.865delC for gRNA-A, c.871delG for gRNA-B) induce frameshifting and premature terminations in the coding sequence, and consequently silence the allele in which they are introduced, other motifs can retain the reading frame (for ex. deletions or insertions of multiples of 3-bp, or splice site indels that would cause in-frame exon 10 skipping). Edits that preserve the reading frame likely produce dominant-negative chains. To account for the functional outcome of the indel edits introduced by Cas9/gRNA-A or –B, we sorted the resulting edits as productive (i.e., desired), neutral, or counter-productive (i.e., undesired). For example, frameshifting indels on the G290R allele were productive, since they silence the dominant-negative allele, whereas frame-preserving indels on either the WT or the G290R allele were counter-productive, since they likely create dominant-negative products. Frameshifting indels on the WT allele were classified as neutral, as they silence this allele but do not cause a dominant-negative product. In patient cells, for gRNA-A, 49.7% of reads were productive, and 25.2% were counter-productive (Figure 1E). In comparison, for gRNA-B, these values were 55.9% and 29.7%, respectively (Figure 1E). Overall, indels introduced by the cleavage of Cas9 in the presence of either gRNA were more frequently productive.

### Addition of intentional mismatches to gRNA-A sequence increases allele specificity

gRNA-A was efficient at inactivating the G290R allele but lacked specificity. To further increase allelic specificity, we deliberately introduced additional base mismatches in the gRNA-A sequence. Each of the re-designed gRNAs now present with two mismatches compared to the WT allele, destabilizing binding and thus editing of the WT allele; meanwhile, the single mismatch compared to the G290R allele should still allow for binding and editing of this allele. We tested three different re-designed gRNAs, in which the deliberate mismatches were introduced in position 2, 3 or 4 of the protospacer (Figure 2A). These gRNAs were nucleofected into the patient and control cell lines, and GFP-enriched cells were analyzed, as described before (Figure S1C). All three re-designed gRNAs were less active than the parental guide at the WT allele; in particular, gRNA-A4a for which only 5.0% of reads included indels in the control sample (Figure 2B). As a trade-off, the attenuated gRNAs were less active at the G290R allele as well (19.7%, 18.5%, and 15.5% of reads were G290R+indels for gRNA-A4a, –A3a, and –A2t, respectively, compared to 32.8% of G290R+indels reads for gRNA-A; Figure 2B). Introduction of the mismatches did not change the editing patterns: deletions, in particular 1-bp deletions, were still the most common edits (Figure S3B,C). Again, the preferential edit was c.865delC (Figure 2C-E). The rate of productive editing was similar for each of the three re-designed gRNAs (69.5%, 65.2%, and 64.7% of reads were productive for gRNA-A4a, –A3a, and –A2t, respectively) and was higher than for the parental gRNA-A (59.5%; Figure 2D).

**Figure 2.**
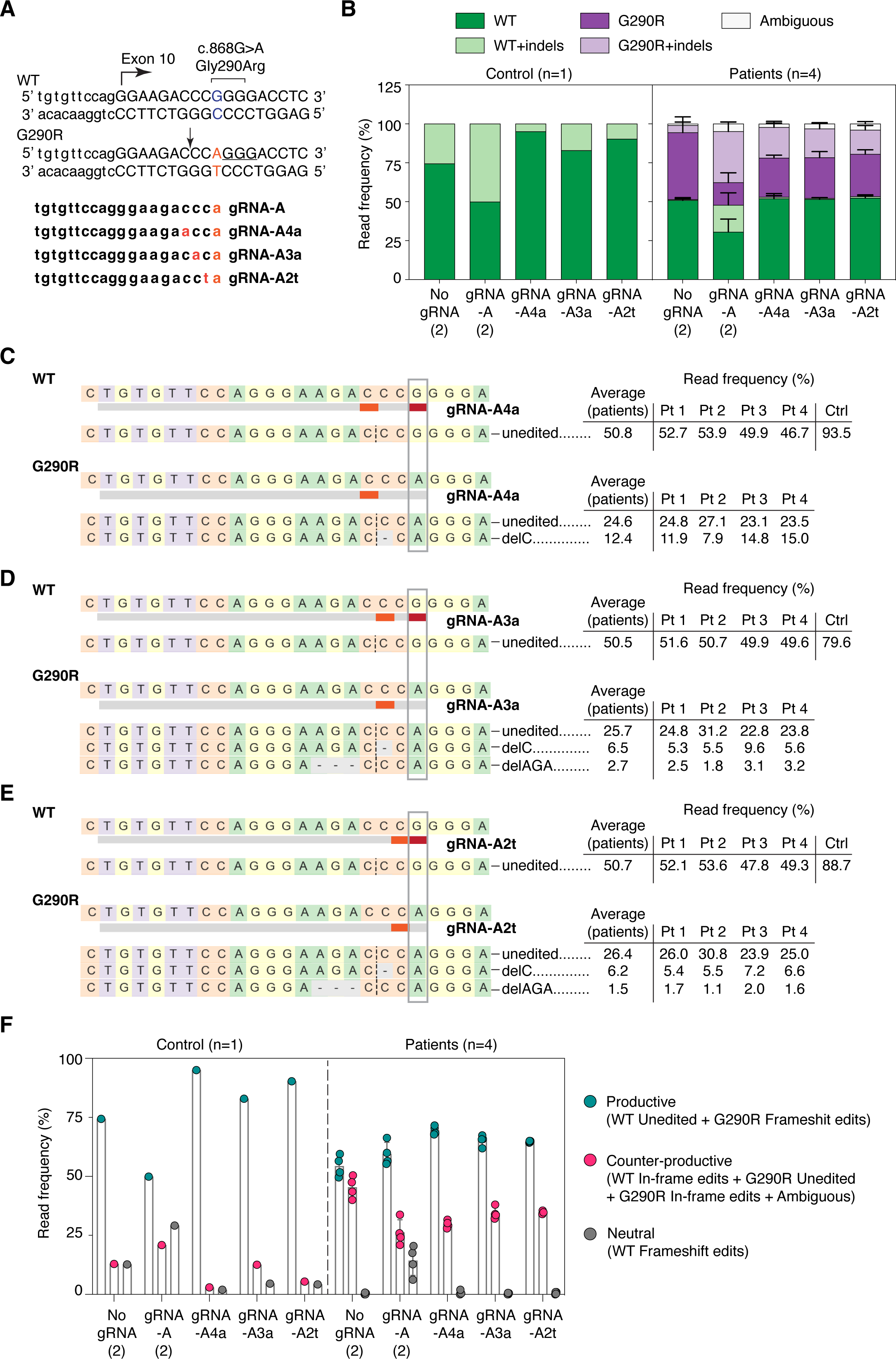
Addition of mismatches to the gRNA sequences increase allele specificity and preserves the repair outcome. **(A)** The gRNA sequence was modified to include an intentional mismatch either at position 4, 3, or 2 of the protospacer. The sequences of the three new attenuated gRNAs, as well as the PAM sequence utilized (underlined) and the Cas9 cleavage site (arrow), are shown. **(B)** Targeted re-sequencing (Illumina Mi-Seq) of the *COL6A1* c.868G>A locus was performed in four patient and one control primary cells after gene editing with gRNA-A, gRNA-A4a, gRNA-A3a, gRNA-A2t, or without gRNA (No gRNA). In this experiment, gRNA-A and No gRNA were replicates from the previous experiment (Figure 1). Sequencing reads were analyzed by Crispresso2 and plotted as in Figure 1. **(C-E)** Alignment of the most frequent reads (reaching a read frequency of at least 1.5% in either the patient or the control samples), for gRNA-A4a **(C)**, gRNA-A3a **(D)** and gRNA-A2t **(E)**. DelC remains the most frequent edit at the G290R allele for all three new re-designed gRNAs. **(F)** Reads were compiled according to the functional outcome, as described in Figure 1. n=4 patient lines ± standard deviation.

### Frameshifting edits effectively inactivate the dominant-negative G290R variant

As a first step to determining the effect of suppressing the G290R allele on collagen VI matrix production, we prepared clonal cell lines following CRISPR/Cas9 editing (Figure S1C). Three clonal cell lines were successfully expanded and analyzed by Sanger sequencing. Clonal cell lines Pt2-gRNA-A and Pt2-gRNA-B acquired frameshifting edits (delC, or delGA;insC) exclusively on the c.868G>A (G290R) allele (Figure 3A). These frameshifting edits are predicted to trigger nonsense-mediated decay, therefore silencing the targeted allele. We confirmed by complementary DNA (cDNA) sequencing that only the c.868G allele was detected (Figure 3B), consistent with absence of expression of the c.868G>A allele. As a result of the G290R allele inactivation, we expected the dominant-negative effect exerted by the mutant collagen VI alpha chain to be alleviated. Indeed, we found that a collagen VI matrix was re-established in Pt2-gRNA-A and Pt2-gRNA-B clonal cells (Figures 3C).

**Figure 3.**
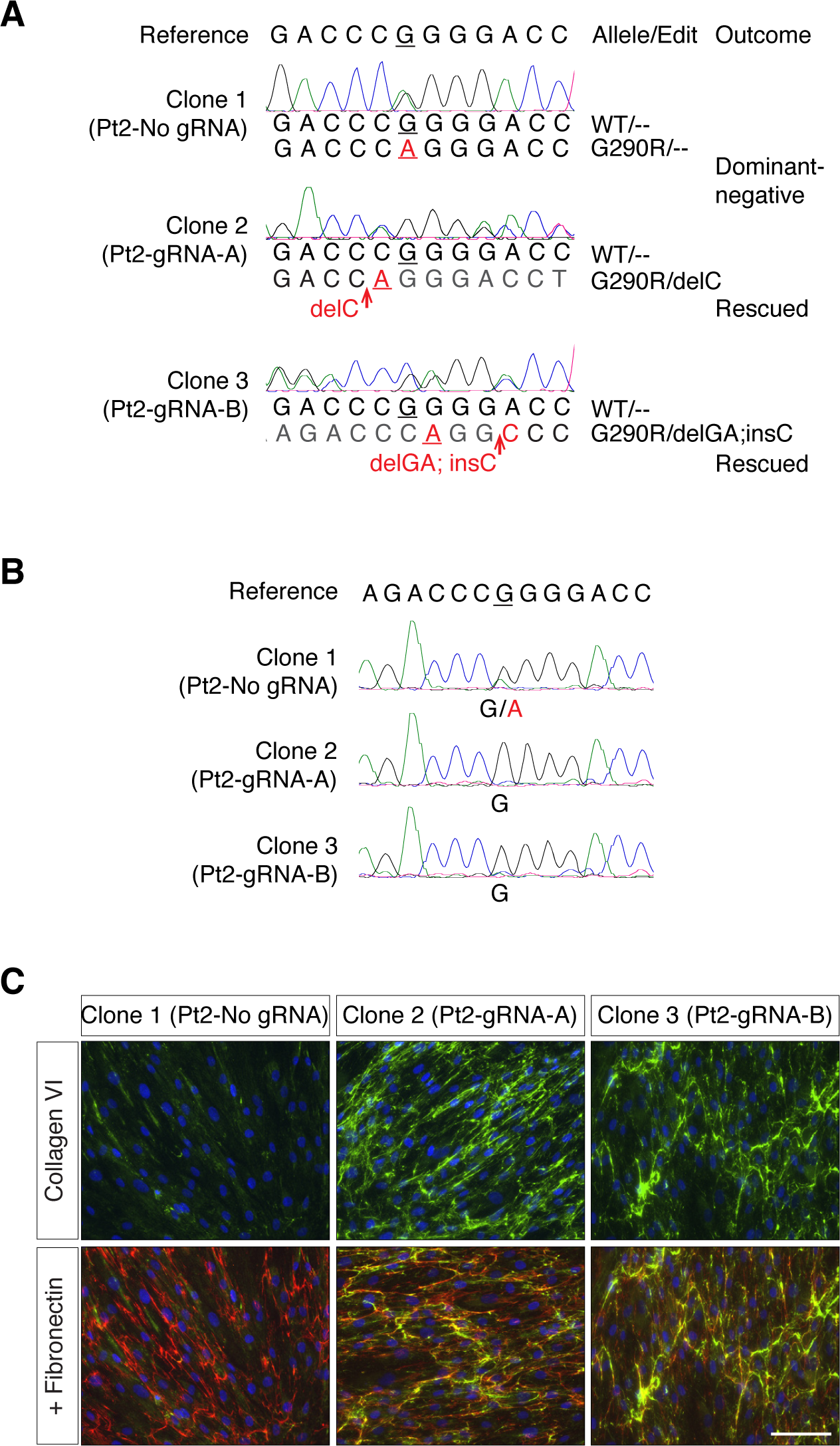
Frameshifting edits at the G290R allele effectively abolish the dominant-negative effect. Patient 2 cells were clonally expanded after gene editing with either gRNA-A, gRNA-B, or without gRNA (No gRNA), and three clones were analyzed. **(A)** Genomic DNA sequencing traces obtained from the three clones (Pt 2-No gRNA, Pt 2-gRNA-A, and Pt 2-gRNA-B) were aligned to the reference sequence. By manually reading either the forward or the reverse traces, the allelic sequences were determined. Clone 2 (Pt2-gRNA-A) includes a delC edit only at the G290R allele, while clone 3 (Pt2-gRNA-B) includes another frameshifting edit (delGA;insC) only at the G290R allele. **(B)** RNA was isolated from the three clonal cell lines to assess expression of *COL6A1*. Complementary DNA (cDNA) sequencing from the clones in (a) shows that clones with G290R allele-specific frameshifting edits exclusively express the WT allele. (**C**) Cells were allowed to reach confluency, after which culture medium was supplemented for three days with ascorbic acid to promote collagen synthesis. Cultured were fixed and immunostained for secreted collagen VI (green) and fibronectin (red). Collagen VI secretion was increased in clonal populations in which the G290R allele is inactivated. Nuclei were identified with DAPI (blue). Bar = 100µm.

### Partial inactivation of the G290R variant improves collagen VI matrix deposition

In a scenario where this CRISPR/Cas9 therapy would be administered systemically, edits could vary from cell to cell. While some cells would have complete abolition of *COL6A1* expression (if both alleles are modified with frameshifting edits) and some would produce a dominant-negative alpha 1 chain (if an in-frame edit is introduced or if there is no editing), a proportion of cells, would, as desired, have a selective disruption of the c.868G>A allele by the introduction of inactivating frameshifts. In this latter case, the dominant-negative effect exerted by the mutant alpha 1 chain would be completely abolished, allowing for the normal alpha 1 chains to form tetramers that secrete, diffuse, and polymerize in the extracellular space, as opposed to the unedited scenario in which every cells carries the dominant-negative acting allele that interferes with collagen VI matrix formation.

To survey the collagen VI matrices produced by mixed populations of edited G290R cells, we treated cells with either of our two initial gRNAs (gRNA-A and –B), or with one of the re-designed gRNAs (gRNA-A4a). We cultured cells in a single dish after GFP enrichment (Figure S1C). Using immunofluorescence and confocal microscopy, we examined the overall collagen VI matrix appearance (Figure 4A), including appearance of the collagen VI microfibrils (Figure 4B), and we quantified the abundance of collagen VI in the cultures (Figure 4C). In the control cell line, only gRNA-A significantly decreased the abundance of the collagen VI matrix, probably due to the number of frameshifting (neutral) edits introduced, but the appearance of the microfibrils did not change. Cultures from patients carrying collagen VI glycine substitutions typically display matrices that are reduced in abundance and that are speckled compared to controls,^9,10^ likely due to increased retention of mutant tetramers and/or reduced adhesion of the secreted tetramers that get washed off during the staining procedure. In addition, while normal matrices show long and continuous microfibrillar structures of collagen VI, mutant glycine matrices usually appear dotty and lack microfibrillar structures. Here, collagen VI abundance in both patient 1 and patient 2 cultures was increased with gRNA-A4a and gRNA-B as opposed to No-gRNA, although it did not reach statistical significance. Interestingly, collagen VI microfibrils in patient cultures appeared more continuous and uninterrupted with all three gRNAs, compared to the No-gRNA control condition. This suggests that reducing the expression of dominant-negative products even in a subpopulation of cells improves overall collagen VI assembly.

**Figure 4.**
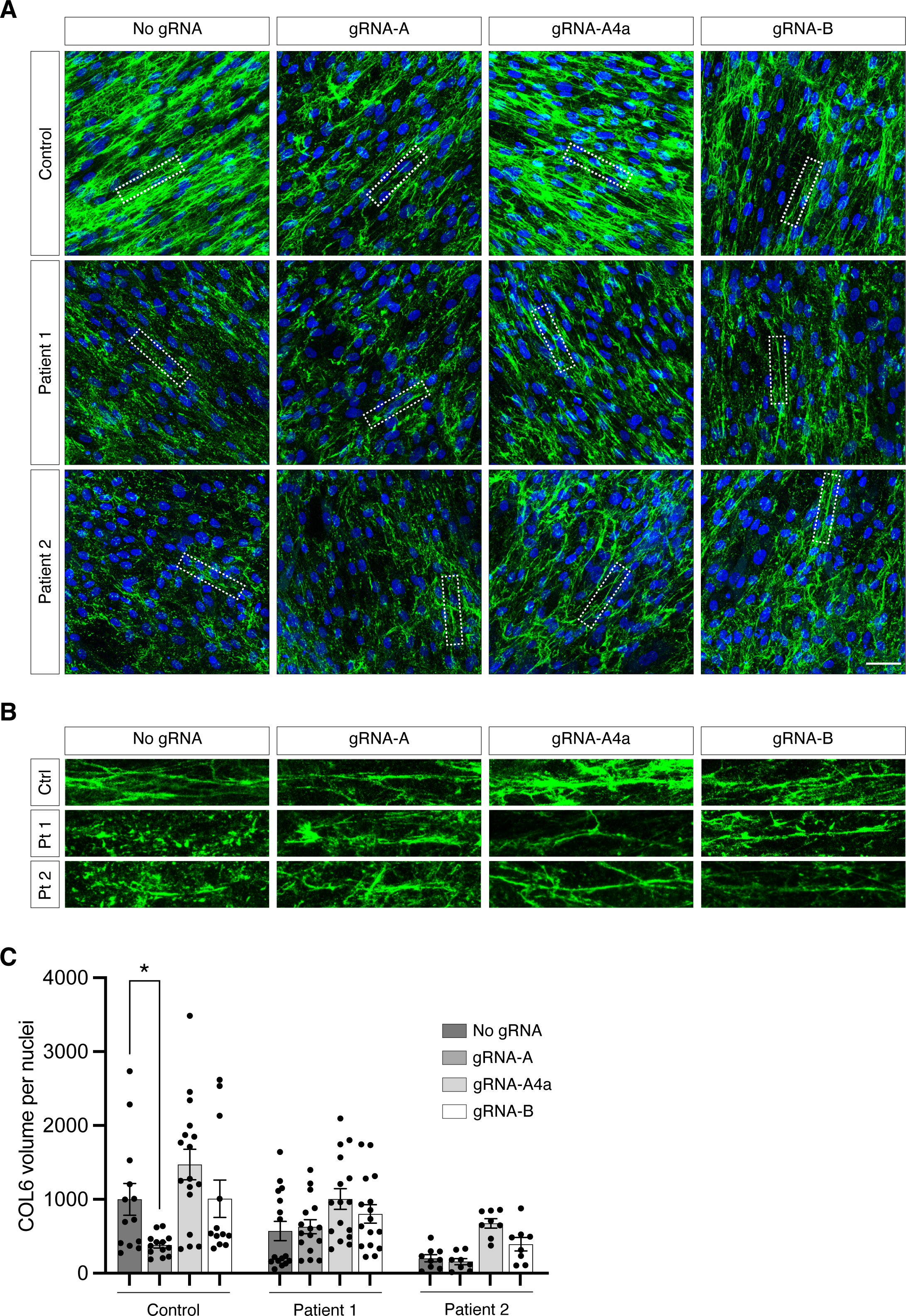
Collagen VI matrix production is increased after gene editing. Cells were collected and expanded in a single dish (mixed cultures) after gene editing with either gRNA-A, gRNA-A4a, gRNA-B or without gRNA (No gRNA). Cultures were immunostained for collagen VI (green) and nuclei (DAPI, blue), and stacks of images were acquired by confocal microscopy. **(A)** Representative merge stacks images showing the abundance of collagen VI secretion in each culture. Bar = 50 µm. **(B)** Enlarged image insets, identified by the dotted rectangles in (A), showing that the structure of the collagen VI microfibrils is dotty in patients’ cultures, but that it is improved after gene editing. **(C)** Quantification of the collagen VI abundance was done by measuring the volume of collagen VI (green fluorescence), normalized to the number of nuclei. Each point represents one quantified field. Four fields were surveyed for each experiment, and experiments were replicated three times. Bars represent average ± standard error of the mean.

Taken together, these data show that random frameshifting edits preferentially introduced at the *COL6A1* c.868G>A (G290R) allele by the action of CRISPR/Cas9 and selective gRNAs can rescue the production of collagen VI.

## Discussion

In this study, we used CRISPR/Cas9 gene editing to introduce frameshifting edits to a recurrent dominant-negative *COL6A1* variant (G290R) to achieve allele-specific gene disruption. Given that variants acting as dominant negative are a frequent cause of COL6-RD and given that harboring a single copy of either of the three *COL6* genes is not associated with a clinical phenotype,^12,13^ allele-specific gene disruption is a suitable approach to treat COL6-RDs. It also bears the advantage that introducing changes at the DNA level, as opposed to the RNA level, can provide permanent inactivation of the dominant allele, allowing edited alleles to be passed on to the progeny of proliferating cells, supporting long-term rescue.

Gene disruption by CRISPR/Cas9 is determined by the resulting DNA repair processes and relies on those to introduce frameshifts into the coding sequence of the targeted gene. Recent findings suggest that end-joining repair profiles following Cas9 cleavage are not random, but for any given gRNA have rather predictable outcomes.^31,32^ In agreement with this notion, our deep-sequencing data showed that the repair profiles were highly similar in four independent primary fibroblast cell lines from patients. Most importantly, the majority of edits observed, both for gRNA-A (and its derivatives) and gRNA-B, were “productive”, i.e. generated the desired frameshifts at the G290R allele. Two single one-bp deletion edits (c.865delC for gRNA-A, –A4a, –A3a, –A2t, and c.871delG for gRNA-B) were predominant and contributed most to the repair profiles. The gRNAs tested in this study effectively introduced frameshifts to the coding sequence and support the use of CRISPR/Cas9-induced gene disruption to inactivate the *COL6A1* G290R variant.

While obtaining productive edits by engaging the cellular DNA repair machinery was successful, achieving allele selectivity was more challenging. In our study, we used a guide-specific approach, in which the variant was in the protospacer rather than in the PAM sequence. The targeting of PAM-creating variants has been shown in several contexts to be the ideal scenario for high allele discrimination, since Cas9 has no detectable activity on the non-targeted allele due to the absence of a PAM sequence.^23,33,34^ However, a pathogenic variant may not necessarily create a PAM site for Cas9, as this was the case for our G290R variant. While in some cases Cas nucleases with different PAM requirements can be considered instead^26,35^, certain positions of the protospacer are thought to also provide sequence specificity, such as the PAM-proximal seed sequence positions 1 to 8 (reviewed in ^36^). Positioning a variant within this region should thus provide allele selectivity. In particular, it was previously reported that an imperfect base pairing rA:dG mismatch at position 1 (like our gRNA-A:protospacer wild-type sequence) nearly abolished cleavage activity.^37^ This was in contrast with our observations, given the suboptimal allele specificity achieved with gRNA-A and gRNA-B in which the variant was in position 1. Additional gRNA design strategies are available to enhance allelic discrimination, such as the utilization of truncated gRNAs,^24,38^ or the introduction of an additional, deliberate, mismatch in the gRNA sequence,^36,39^ which is the strategy we selected here. In our experimental design, introduction of a mismatch at a 3-nucleotide distance of the variant (gRNA-A4A) destabilized sufficiently hybridization to the WT allele to almost abrogate cutting activity completely in a control cell line. This simple design strategy adds to the design options for precision CRISPR/Cas therapeutics and could be combined with the use of engineered Cas9 with greater fidelity,^40–42^ or with the use of alternate Cas nucleases.^43,44^

A recent study applied this allele-specific gene disruption strategy to a different recurrent *COL6A1* glycine substitution (G293R) and reported high allele-specificity and productive introduction of frameshifting edits.^45^ By incorporating the intentional mismatches in the gRNA design, we achieved allele specificity to levels comparable to the López-Márquez study for our target as well. One factor that could have contributed to the higher specificity observed by López-Márquez et al. using an unaltered gRNA is the position of the variants within the protospacer, which were located at either position 2 or 8.^45^ Factors other than the primary sequence, however, may also account for the different specificity levels we and López-Márquez et al. observed, such as the cellular concentration of Cas9/gRNA and the time of exposure to Cas9 nuclease.^37^ We used plasmid-based expression of Cas9 and gRNAs for 48h, that ‘mimics’ a viral delivery; López-Márquez et al., instead, delivered CRISPR components as ribonucleoprotein complexes that reflect best a nanoparticle-based delivery.^45^ In these current study designs, however, it is not possible to determine the contribution of each of these factors to the levels of allele specificity observed. Similarly to the López-Márquez study, we observed productive NHEJ repair outcomes and improvement of collagen VI matrix production, further supporting this CRISPR/Cas9-induced gene disruption strategy. The two studies together demonstrate that CRISPR/Cas9 approaches can be tailored to different recurrent dominant disease-causing glycine variants in the *COL6A1* gene, and thus likely also to other such disease-causing variants in the three *COL6* genes.

Dominant variants on either of the three *COL6* genes, such as glycine substitutions located at the N-termini of the triple helical domains, frequently have profound dominant-negative effects on collagen VI assembly when they are “assembly competent”, i.e. when they can be carried forward into to the final secreted product, the multimers of α1(VI), α2(VI) and α3(VI) protein chains that constitute the collagen VI extracellular matrix. Of note, patients who carry such variants in somatic mosaicism show considerably milder phenotypes.^46,47^ Hence, even a subpopulation of cells capable of depositing a normal collagen VI matrix, as an *in vivo* CRISPR/Cas9 therapy would likely produce, has the potential to ameliorate the disease. Additionally, as noted before, every corrected cell is expected to generate an expanding progeny of corrected cells. Here we show that clonally expanded correctly edited cells show a normalized matrix deposition, but also realize that the potential effect *in vivo* is best modeled in our experiment where we kept primary cells as a mixed culture after editing has been applied. Testing this experimental scenario we found that the overall extracellular microfibrillar structure of collagen VI in the cultures was also improved. While there are no established assays to verify functional recovery of this matrix in culture, the qualitative improvement we report is encouraging and consistent with the observation of the milder phenotype observed in the individuals somatically mosaic for dominant-negative *COL6* variants. Additional experimentation is needed to determine what threshold of corrected cells is necessary to attain a restored and functional matrix, but these data provide an excellent rationale for allele-specific gene disruption of collagen VI genes.

Collagen VI in muscle is mainly expressed by muscle interstitial fibroblasts.^3,4^ As a cellular model, we used here primary fibroblastic cells derived from normal or patient skin. Skin-derived fibroblasts are capable of producing and depositing collagen VI efficiently and are well established to reflect the abnormal collagen VI deposition in wide variety of COL6-RD patients.^30^ However, given that the genomic and epigenetic context influences Cas9 activity, working with a relevant cell type is essential, in particular for the survey of genomic off-targets, which we did not evaluate systematically in the current study.

CRISPR/Cas9 is a powerful gene editing tool that can mediate targeted gene disruption. Two recent phase I clinical studies made use of this experimental design, in which Cas9 and gRNAs were administered *ex vivo* to transfusion-dependent β-thalassemia, and sickle cell disease patients,^48^ and *in vivo* to transthyrethin amyloidosis patients.^49^ In these trials, expression levels of the targeted genes were strongly reduced; however, the gRNAs were designed to target both alleles. Modifying the system to achieve allele-specific gene disruption would expand the application to additional conditions, including conditions caused by dominant and dominant-negative variants^19^ that occur in haplo-sufficient genes. Our study focused on a variant-specific approach; however, in combination with the Lopez et al. study, it provided a strong proof-of-principle for the application of this approach to other dominant *COL6* variants. Our study also supports mismatch gRNA design as an additional tool to the precision engineering of allele-specific gRNAs. To make this approach into a more universal tool for the COL6-RD patient community, an interesting strategy would be to develop a repertoire of allele-specific gRNAs for each allele of common single nucleotide polymorphisms, and to target those polymorphisms as proxy for pathogenic variants located in *cis* to the polymorphisms.^50^

## Materials and Methods

### Guide RNA Design

Possible guide RNA (gRNA) designs were analyzed *in silico* using the Broad Institute’s sgRNA Designer (http://www.broadinstitute.org/rnai/public/analysis-tools/sgrna-design), now CRISPick (https://portals.broadinstitute.org/gppx/crispick/public)^51,52^ and the Massachusetts Institute of technology’s former CRISPR Design (http://www.crispr.mit.edu)^37^ tools. Guide RNAs overlapping the pathogenic variant site were listed and compared for their on-target score, number of off-targets, and the distance of the mutation site to the PAM sequence (Figure S1A,B). The number of off-target matches reported includes coding regions and non-coding regions of coding genes as well as cutting frequency determination (CFD) scores between 0.2 and 1.0, inclusively. Given the desire for allele discrimination, the two guide sequences chosen were those for which the mutation site was in the seed sequence proximal to the PAM sequence and that scored highest for each design tool.

### Cloning

Plasmid pSpCas9(BB)-2A-GFP (PX458) was obtained from Addgene #48138 (Watertown, MA) courtesy of Dr. Feng Zhang, and selected guide sequences were cloned according to the protocol described in Ran et al.^53^ Following cloning, plasmid DNA was isolated using an endotoxin-free maxiprep kit (Qiagen, Germantown, MD). Plasmids were verified by Sanger sequencing (Genewiz, South Plainfield, NJ).

### Subjects

Skin biospies were obtained based on standard operation procedures from subjects recruited at the National Institutes of Health (NIH) through the Pediatric Neuromuscular Clinic (12-N-0095, approved by the Institutional Review Board of the National Institute of Neurological Disorders and Stroke (NINDS), NIH).

### Cell Culture and Nucleofection of Cas9/guideRNA Plasmids

Normal control and patient dermal fibroblast cell lines were established from skin biopsies. Cells were maintained in Dulbecco’s modified Eagle medium (DMEM, Gibco/ThermoFisher Scientific, Waltham, MA) with 10% fetal bovine serum (FBS) (Corning/ ThermoFisher Sci) and 1% penicillin/streptomycin (P/S) (Gibco/ThermoFisher Sci) in 5% CO_2_ at 37°C. Prior to conducting experiments, cells were tested for mycoplasma contamination (MycoAlert mycoplasma detection kit, Lonza, Basel, Switzerland) and treated if necessary (MycoZap Reagent, Lonza). At the time of nucleofection, 2-4 x 10^5^ cells were resuspended in 100uL of P2 medium (Lonza) containing 10 µg of Cas9 or Cas9/guide plasmidic DNA and transferred to a cuvette (Lonza). Cells were nucleofected with the DT-130 program using the 4D-Nucleofector system with Core and X unit (Lonza). Nucleofected cells were incubated for 10min at room temperature before being transferred to a 6-well plate containing pre-warmed and equilibrated medium (DMEM + 10% FBS). After 48 hours, cells were trypsinized, washed twice with phosphate buffered saline (PBS), and resuspended in 500 uL of cell sorting medium (145 mM NaCl, 5 mM KCl, 1.8 mM CaCl_2_, 0.8 mM MgCl_2_, 10 mM Hepes, 10 mM Glucose) containing 1µg/mL of 4′,6-Diamidino-2-phenylindole dihydrochloride (DAPI) (Sigma-Aldrich, St. Louis, MO).

### Cell Sorting and Cell Culture

Cells were sorted at the NINDS Flow and Imaging Cytometry Core Facility on the MoFlo Astrios Cell Sorter (Beckman Coulter Life Sciences, Indianapolis, IN). The strategy for gating was to first exclude dead cells using 1µg/mL DAPI staining, and then sort on viable DAPI-negative cells that also expressed GFP. Threshold for GFP expression was established using untransfected cells as controls. GFP^+^ cells were either sorted in complete culture medium for cell culture and DNA extractions (DMEM + 10% FBS + 1% P/S + 1% amphotericin B), or directly in Trizol LS (Invitrogen/Thermo Sci) for RNA extractions.

For cell culture, cells were pelleted and washed once in culture medium, and then were either resuspended in complete medium and plated in a single dish, or they were resuspended in conditioned medium and plated by serial dilution in 96-well plates. Conditioned medium was obtained by collecting medium from wild-type primary fibroblasts that were cultured for at least two days to confluency, and by filtering the conditioned medium with 0.2 µm filters. In the 96-well plates, wells in which a single cell had been seeded were flagged for observation. Once cells reached confluency in the flagged wells, they were successively expanded in larger wells until the population size was large enough to perform assessments.

### DNA/RNA Isolation, Amplification, and Sanger Sequencing

DNA was isolated from fibroblast cells using the Puregene Kit (Qiagen) according to instructions. DNA was amplified using the 2X KAPA Taq ReadyMix PCR Kit (KAPA Biosystems, now Roche Sequencing, Indianapolis, IN) for 35 cycles using an annealing temperature of 64°C. Primers for this reaction were as followed: F, 5’-cacactgcctgttccttgtg –3’, and R 5’-gtcgagcctcactcaccttc-3’. PCR products were sent for purification and Sanger sequencing (Genewiz), and sequencing reads were aligned to the reference sequence using DNASTAR Lasergene (DNASTAR, Madison, WI). Sequence chromas were read manually to analyze the composition of edited alleles at the c.868G>A location.

Total RNA was isolated from fibroblast cells with Trizol (Invitrogen/ThermoFisherSci) according to manufacturer’s instructions. 500 ng of RNA were treated with recombinant DNaseI using the DNA-free DNA Removal Kit (Ambion/ThermoFisher). The DNA-free RNA sample was then used for reverse transcription, using the SuperScript III Reverse Transcriptase (Invitrogen/ThermoFisher) and random primers. Amplification was performed using the Advantage 2 Polymerase Mix (Clonetech, now Takara Bio USA, Mountain View, CA) and the following primers: F, 5’-ccatcgtggacatgatcaaa-3’, and R 5’-ccctcgtctccagatggtc-3’. Sanger sequencing was performed and analyzed as described above.

### Targeted Re-Sequencing

Following cell sorting, cells were pelleted by centrifugation at 10,000xg for one minute and washed once with PBS. DNA was isolated as described above. DNA library for targeted re-sequencing was prepared using the following procedure. A first PCR reaction was prepared from genomic DNA using the KAPA HiFi HotStart PCR Kit (KAPA Biosystems/Roche Sequencing) in a total reaction volume of 20 µL, with an annealing temperature of 68°C for 15 cycles (except for the three Patient 3 samples for which 25 cycles were used at this step because of low DNA yield). The forward primers for this reaction (5’-ctccttggcccaaatcctat-3’) were each extended in 5’ with a nucleotide stagger, a barcode unique to each sample, and the Illumina forward extension (5’-acactctttccctacacgacgctcttccgatct-3’), while the reverse primers (5’-agagaccagctccgaggtc-3’) were also extended with a nucleotide stagger, a barcode unique to each sample, and the Illumina reverse extension (5’-gtgactggagttcagacgtgtgctcttccgatct-3’). A second round of amplification was performed, in duplicate, using 8 µL of PCR products from the first round, using the same conditions as above, except for the number of cycles which was increased to 25. The primers used for that second round were the Illumina adapter primers forward (5’-aatgatacggcgaccaccgagatctacactctttccctacacgac-3’) and reverse (5’-caagcagaagacggcatacgagatgtgactggagttcagacgtgt-3’). Reaction duplicates were pooled and purified with the QIAquick PCR Purification Kit (Qiagen). Products were sent for targeted re-sequencing on the Illumina MiSeq platform at the NIH Intramural Sequencing Center (https://nisc.nih.gov/index.htm). For the second series of targeted re-sequencing experiments (Figure 2), the same protocol was applied, with the exception of the gene-specific reverse primer in the first round of amplification that was modified (5’-tagtgctgtgcaaggctgag-3’) to generate a shorter product.

Paired-End sequencing reads were analyzed with CRISPResso2 (http://crispresso.pinellolab.partners.org)^54^ using the default settings, with the exception of the quantification window size which was set to 4bp. Reads were classified as unedited or containing indels. Reads that could not be called as either WT or G290R, because the indel encompassed the c.868 nucleotide site, were classified as ambiguous.

### Immunostaining

Fibroblasts (0.8 x 10^4^) were seeded into 8-chamber tissue culture slides (Corning/ThermoFisher Sci). Culture medium was replaced the following day to supplement with 50 µg/mL of L-ascorbic acid (Wako Chemicals USA, Richmond, VA). After two days, medium was once again replaced to maintain a sufficient supply of L-ascorbic acid. After a total of three to four days of L-ascorbic acid treatment, cells were washed 1x in PBS and then fixed for 10 minutes at room temperature in 4% paraformaldehyde (Electron Microscopy Sciences, Hatfield, PA). Matrix immunostaining was performed using the same procedure and antibodies as described previously.^55^

### Microscopy

Epifluorescence images were acquired on an inverted Nikon Eclipse Ti microscope (Nikon Instruments, Melville, NY) equipped with a sCMOS pco.edge 4.2 LT camera (Excelitas PCO GmbH, Kelheim, Germany). Confocal images were acquired using a TCS SP5 II system (Leica Microsystems, Buffalo Grove, IL), with 40X or 63X objectives. A thickness of 0.508 µm was obtained, and z-stacks were acquired using 0.5 µm-sized steps. Collagen VI matrix deposition was quantified according to methods set forth in Bolduc et al.^17^

## Data Availability Statement

Targeted re-sequencing data from this study have been submitted to the NCBI Sequence Read Archive (BioProject ID PRJNA1023208).

## Acknowledgements

We thank Dragan Maric at the NINDS Flow and Imaging Cytometry Core Facility, as well as Alice Young and the NIH Intramural Sequencing Center staff. We thank Fady Guirguis and Janelle Hauserman for their input. We thank Feng Zhang for providing the Cas9 plasmid (Addgene #48138). This work was supported by the Division of Intramural Research of the NIH, NINDS (1ZIANS003129). The content is solely the responsibility of the author(s) and does not necessarily represent the official views of the National Institutes of Health.

## Author contributions

Conceptualization, V.B., K.S. and C.G.B.; Methodology, K.S., V.B. and A.S.; Investigation, V.B., K.S., E.E., A.B., G.S.C. and P.U.; Data curation, K.J., E.E. and P.U.; Software: E.E., P.U.; Writing – Original draft, V.B.; Writing – Review and editing, V.B., K.S., E.E., A.B., G.S.C., A.S., P.U., C.G.B.; Visualization, V.B., K.S., E.E. and A.B.; Supervision: V.B. and C.G.B.; Funding, C.G.B.

