## Supplementary Figures for "Allele-specific CRISPR/Cas9 editing inactivates a single nucleotide variant associated with collagen VI muscular dystrophy"

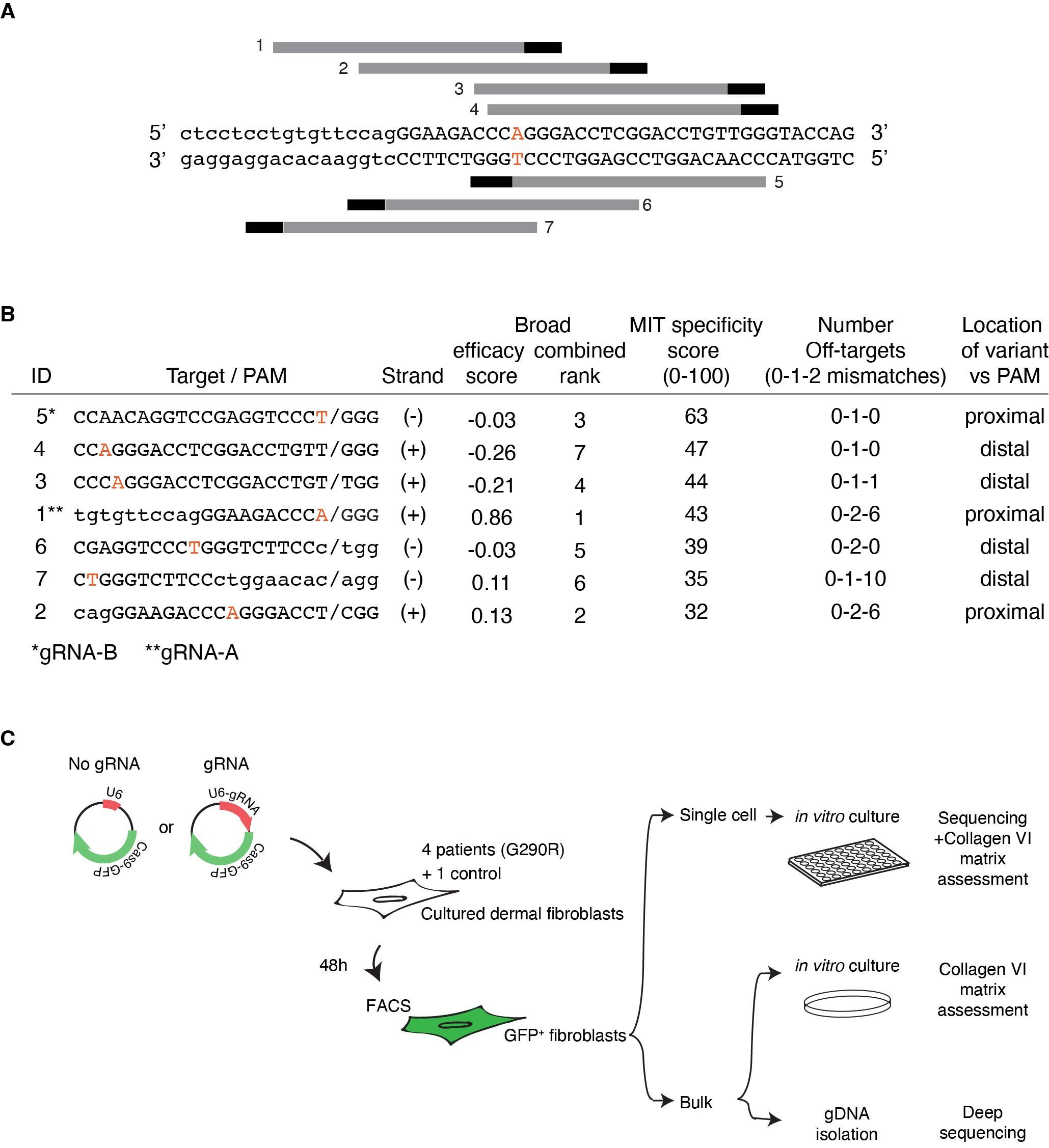

**Figure S1. Strategy for allele-specific gene editing of the *COL6A1* c.868G>A (G290R) variant.**

**(A)** The localization of potential allele-specific gRNAs for the c.868G>A variant is illustrated along the *COL6A1* genomic sequence. gRNAs hybridizing to the negative (-) strand are shown below, while gRNAs hybridizing to the positive (+) strand are shown above the sequence. **(B)** The sequences of all potential gRNAs are listed, together with the sequences of the adjacent PAM sites. The presence of the mutation within a 10-nt distance of the PAM site was considered proximal. ID 1 was chosen as gRNA-A and ID 5 was chosen as gRNA-B. **(C)** Constructs expressing a Cas9-GFP fusion protein, in combination with either of the gRNAs, or without gRNA (no gRNA), were each nucleofected into human primary dermal fibroblasts (4 patients and one unaffected control). After 48 hours, cells were sorted for GFP expression (GFP^+^). GFP^+^ cells were plated with a serial dilution method to isolate single clones for sequencing and collagen VI matrix assessment on pure cell populations, or GFP^+^ cells were plated and expanded in a single dish (bulk) for collagen VI matrix assessment on mixed populations. Alternatively, genomic DNA extraction was performed on GFP^+^ cells for Illumina Mi-Seq targeted re-sequencing and analysis.

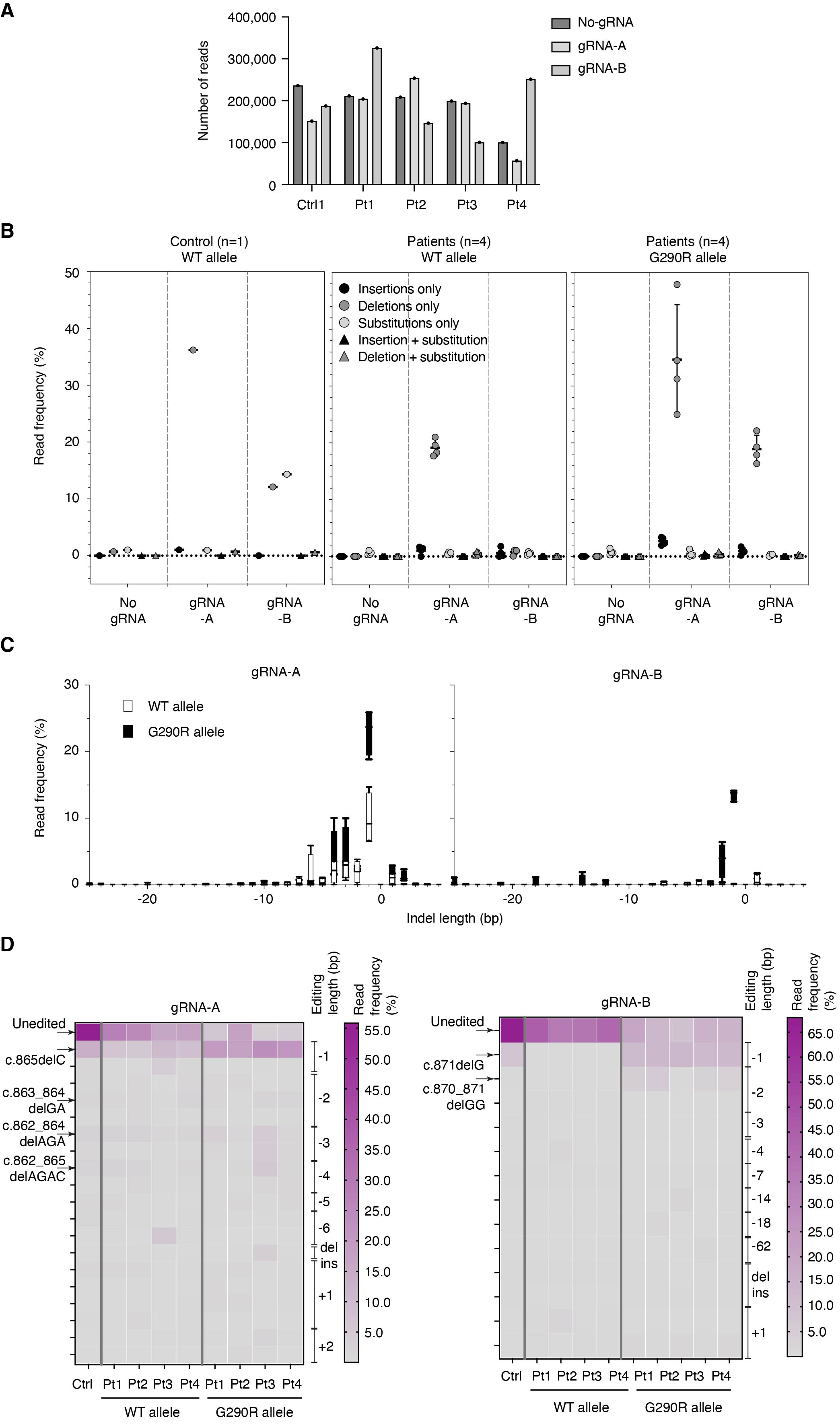

**Figure S2. Targeted re-sequencing analysis of the repair outcomes after gene editing with gRNA-A or gRNA-B.**

Four patient and one control primary cells were co-transfected with Cas9-GFP and with either gRNA-A, gRNA-B, or without gRNA (No gRNA). GFP^+^ cells were collected after 48h, and genomic DNA was subjected to targeted re-sequencing (Illumina Mi-Seq) of the *COL6A1* c.868G>A locus. Sequencing reads were analyzed by Crispresso2. **(A)** Total number of sequencing reads, per sample. The total number of reads was subsequently used as the denominator to calculate the read frequency in each sample. **(B)** Total read frequencies per indel/edit type (deletions, insertions, substitutions, or combination thereof) and per allele type (WT or G290R). Bars represent average ± standard deviation. **(C)** Total read frequencies per indel length, and per allele type, reported as box and whisker plots, for gRNA-A (left) and gRNA-B (right). On the x axes, the negative scale represents length of deletions, while the positive site represents length of insertions. **(D)** Heatmaps of the top 20 motifs for their total read frequencies, identified at the G290R allele in patient samples, and displayed by individual.

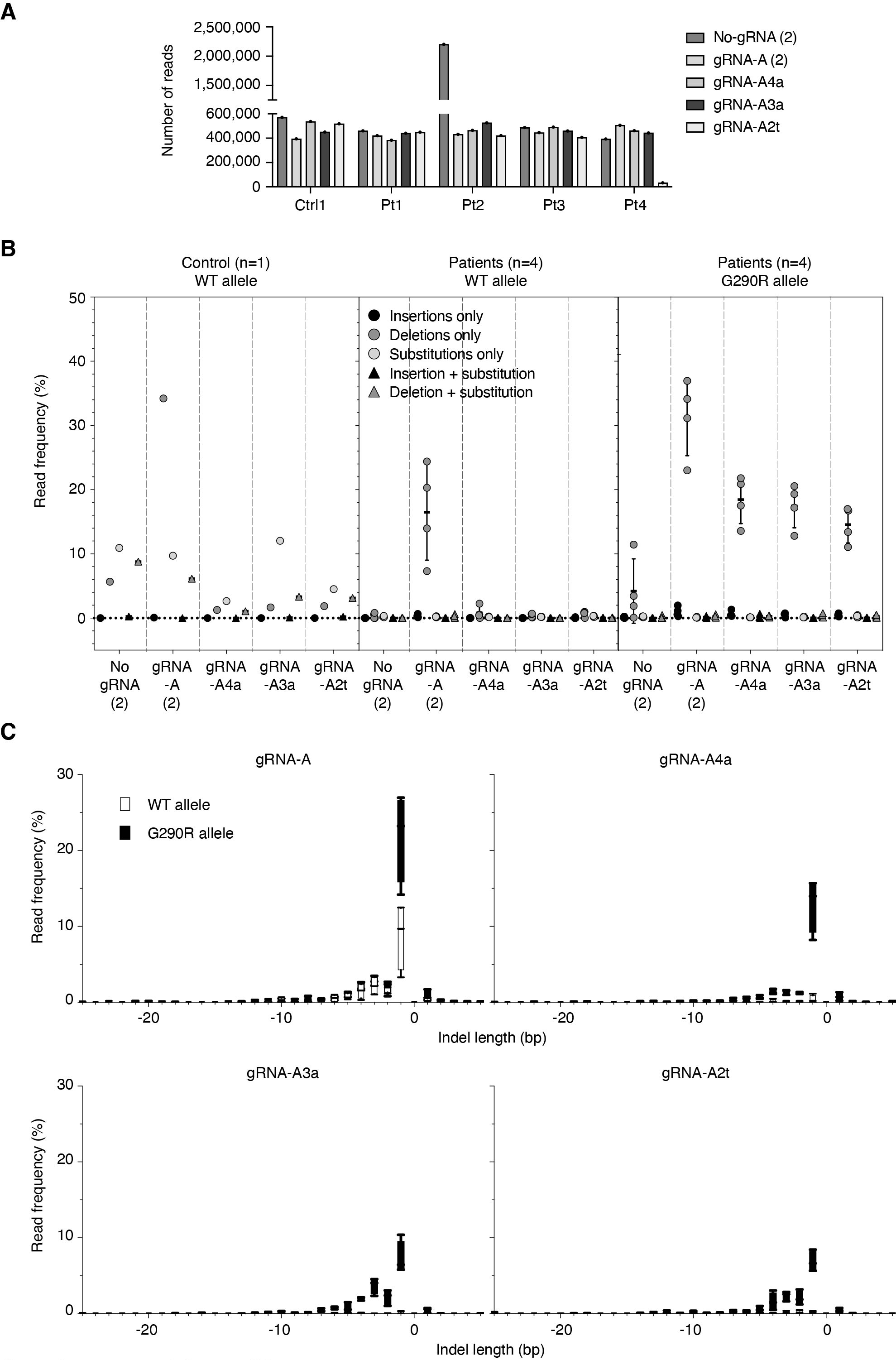

**Figure S3. Targeted re-sequencing analysis of the repair outcomes after gene editing with gRNA-A4A, gRNA-A3A or gRNA-A2T.**

Four patient and one control primary cells were co-transfected with Cas9-GFP and with either gRNA-A, gRNA-A4a, gRNA-A3a, gRNA-A2t, or without gRNA (No gRNA). GFP^+^ cells were collected after 48h, and genomic DNA was subjected to targeted re-sequencing (Illumina Mi-Seq) of the *COL6A1* c.868G>A locus. Sequencing reads were analyzed by Crispresso2. Note that gRNA-A and No gRNA in this experiment are replicates from the previous experiment. **(A)** Total number of sequencing reads, per sample. The total number of reads was subsequently used as the denominator to calculate the read frequency in each sample. **(B)** Total read frequencies per indel/edit type (deletions, insertions, substitutions, or combination thereof) and per allele type (WT or G290R). Bars represent average ± standard deviation. **(C)** Total read frequencies per indel length, and per allele type, reported as box and whisker plots, for each of the gRNA. On the x axes, the negative scale represents length of deletions, while the positive site represents length of insertions. **(D)** Heatmaps of the top 20 motifs for their total read frequencies, identified at the G290R allele in patient samples, and displayed by individual.
